# Breaking the Burst: Unveiling Mechanisms Behind Fragmented Network Bursts in Patient-derived Neurons

**DOI:** 10.1101/2024.02.28.582501

**Authors:** Nina Doorn, Eva J.H.F. Voogd, Marloes R. Levers, Michel J.A.M. van Putten, Monica Frega

## Abstract

Fragmented network bursts (NBs) are observed as a phenotypic driver in many patient-derived neuronal networks on multi-electrode arrays (MEAs), but the pathophysiological mechanisms underlying this phenomenon are unknown. Here, we used our previously developed biophysically detailed *in silico* model to investigate these mechanisms. Fragmentation of NBs in our model simulations occurred only when the level of short-term synaptic depression (STD) was enhanced, suggesting that STD is a key player. Experimental validation with Dynasore, an STD enhancer, induced fragmented NBs in healthy neuronal networks *in vitro*. Additionally, we showed that strong asynchronous neurotransmitter release, NMDA currents, or short-term facilitation (STF) can support the emergence of multiple fragments in NBs by producing excitation that persists after high-frequency firing stops. Our results provide important insights into disease mechanisms and potential pharmaceutical targets for neurological disorders modeled using hiPSC-derived neurons.

## Introduction

Human induced pluripotent stem cell (hiPSC)-derived neurons have emerged as an effective platform for drug screening and modeling of neurological disorders *in vitro*. When plated on multi-electrode arrays (MEAs), these neurons form functionally connected and spontaneously active networks [1]. This activity self-organizes into network bursts (NBs), which are drastic transient increases in spiking frequency occurring synchronously throughout the network. The properties of these NBs are often used for phenotypic characterization as they correlate with specific disease states [2, 3, 4, 5, 6]. Several of these genotype/phenotype correlations have been established by characterizing NBs, providing insight into the pathophysiological mechanisms underlying the neuronal network phenotype [7, 3, 4, 5, 6, 8, 9].

While in healthy neuronal networks NBs show a monotonous decrease in spiking frequency back to baseline activity, NBs in patient-derived neuronal networks may express prominent fluctuations in the spiking frequency within this period, resulting in *fragmented* NBs. In research using neuronal networks derived from hiPSCs of patients with Dravet Syndrome (DS), GEFS+, and Febrile Seizures with mutations in *SCN1A*, fragmented NBs were the main phenotypic driver [2]. Anti-epileptic drugs (AEDs) that were effective in those patients reduced the number of fragments per NB in their neuronal networks, while AEDs that exacerbated the clinical phenotype increased it. Similarly, in a model for Kabuki Syndrome, a multisystem neurodevelopmental disorder (NDD), fragmented NBs were one of the main signs of altered network organization compared to healthy networks [10]. Fragments in NBs were also visible in other NDD models such as Kleefstra Syndrome (KS) [5] and Rett syndrome (RTT) [11], and were suppressed with either NMDA receptor-, or asynchronous neurotransmitter release blockers. However, how these processes contribute to the emergence of fragments remains unknown. Uncovering the complete pathways leading to the fragmentation of NBs could help elucidate disease mechanisms at play in patient-derived neuronal networks, and provide possible pharmaceutical targets.

Computational models can provide insight into the mechanisms underlying specific electrophysiological behavior. We previously developed a biophysically detailed computational model of hiPSC-derived neuronal networks on MEAs, that can faithfully simulate the pattern of activity of healthy and patient-derived neuronal networks [7]. Here, we use this model to investigate the possible mechanisms underlying fragmented NBs.

Fragmentation of NBs in our model simulations occurred only when the level of short-term synaptic depression (STD) was substantial, suggesting this is a key player. We validated our hypothesis by increasing STD in healthy neuronal networks *in vitro* with Dynasore, which resulted in the emergence of fragmented NBs. Furthermore, we showed that enhanced STD in combination with persistent excitation supports the generation of multiple fragments in NBs. Our study provides crucial insights into disease mechanisms at play in neuronal network models of neurological disorders.

## Results

### Fragmented NBs as a common phenotype in patient-derived neuronal networks

Fragmented NBs are observed in multiple patient-derived neuronal networks grown on MEAs. To illuminate the extent of this phenotype, we gathered MEA measurements of such networks. Specifically, we re-analyzed data from networks derived from patients with GEFS+ and DS (mutation in *SCN1A*) [2], and KS (mutation in *EHMT1*) [5], as well as the corresponding healthy control networks. In all cases, patient-derived stem cells were differentiated into excitatory neurons through forced *Ngn2* overexpression (Figure 1A). The activity of all neuronal networks consisted of random spiking and NBs. Compared to controls, patient-derived neuronal networks showed altered NB characteristics, which were disease-specific (i.e., more or less frequent, shorter or longer NBs, Figure 1B). Upon examination of recordings from a single electrode during NBs, it becomes apparent that these phenotypes have a shared characteristic, namely the presence of fragments, resulting from prominent fluctuations in the spiking frequency within the NB (Figure 1C, top panels). The number and ‘shapes’ of these fragments vary widely between these disorders. To quantify the number of fragments, we detected the local maxima in the smoothed network firing rate for each NB (see Methods for details), indicated with the colored dots in Figure 1C, bottom panels. We observed significantly increased numbers of fragments in patient-derived networks compared to control (Figure 1D). Thus, Fragmented NBs were a common phenotype of patient-derived neuronal networks on MEAs.

**Figure 1:**
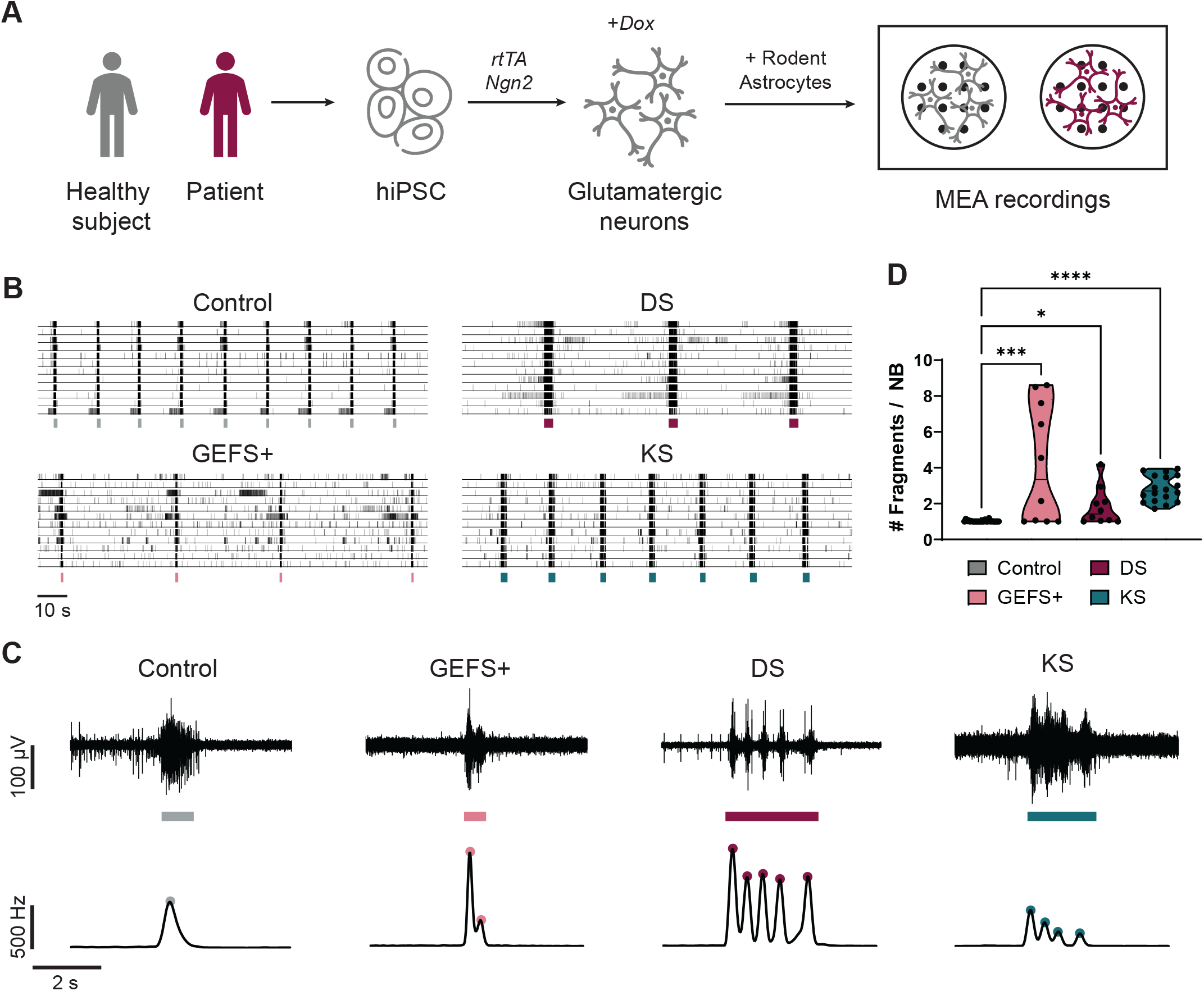
Fragmented network bursts in patient-derived neuronal networks on MEA. **A)** Schematic overview of the protocol used by [2, 5] to differentiate hiPSCs into neuronal networks on multi-electrode arrays (MEAs). hiPSCs were obtained by reprogramming somatic cells of healthy subjects and patients. Excitatory neurons were generated through doxycycline (Dox) inducible overexpression of Neurogenin2 (*Ngn2*). At Day *In Vitro* (DIV) 2, E18 rat astrocytes were added in a 1:1 ratio. Activity was recorded at DIV 35 with MEA. **B)** Representative rasterplots of spontaneous activity of healthy control neuronal networks (Control) and networks derived from patients with GEFS+, Dravet Syndrome (DS), and Kleefstra Syndrome (KS) recorded by [2, 5]. Detected network bursts (NBs) are indicated by the colored bars below. **C)** Top: representative example voltage traces recorded at one electrode during an NB detection (bar below). Bottom: representative network firing rate traces during the same NB and detected fragments (colored dots). **D)** Quantification of the average number of fragments per NB for Control (n=26), GEFS+ (n=10), DS (n=11), and KS (n=18) networks. * P*<*0.05, *** P*<*0.001, **** P*<*0.0001, Kruskal-Wallis test with Dunn’s multiple comparisons test was performed between groups.

### Our biophysical models replicate fragmented NBs with increased short-term synaptic depression

Next, we aimed to investigate the mechanisms underlying the appearance of fragmented NBs in patient-derived neuronal networks. For this, we employed our previously developed computational model of hiPSC-derived excitatory neuronal networks on MEA [7], which has been shown to successfully identify the effect of specific cellular changes on the network activity.

Our biophysical model (Figure 2A) consists of heterogeneous Hodgkin-Huxley (HH) neurons, connected through synapses with AMPA- and NMDA-receptors (AMPAr and NMDAr), each with varying strengths and delays. The neurons contain voltage-gated potassium and sodium channels as well as slow-afterhyperpolarizing (sAHP) channels causing spike-frequency adaptation. Moreover, the synapses undergo short-term synaptic depression (STD): the amplitude of the excitatory postsynaptic current (EPSC) is depressed with every subsequent presynaptic spike and recovers quickly in the absence of spikes. The activity of the network is recorded by virtual electrodes, mimicking the experimental MEA recordings. The simulated activity of this model accurately recapitulated the NB characteristics of control neuronal networks at the single electrode level (Figure 2B “Basal” and Figure 1C “Control”).

**Figure 2:**
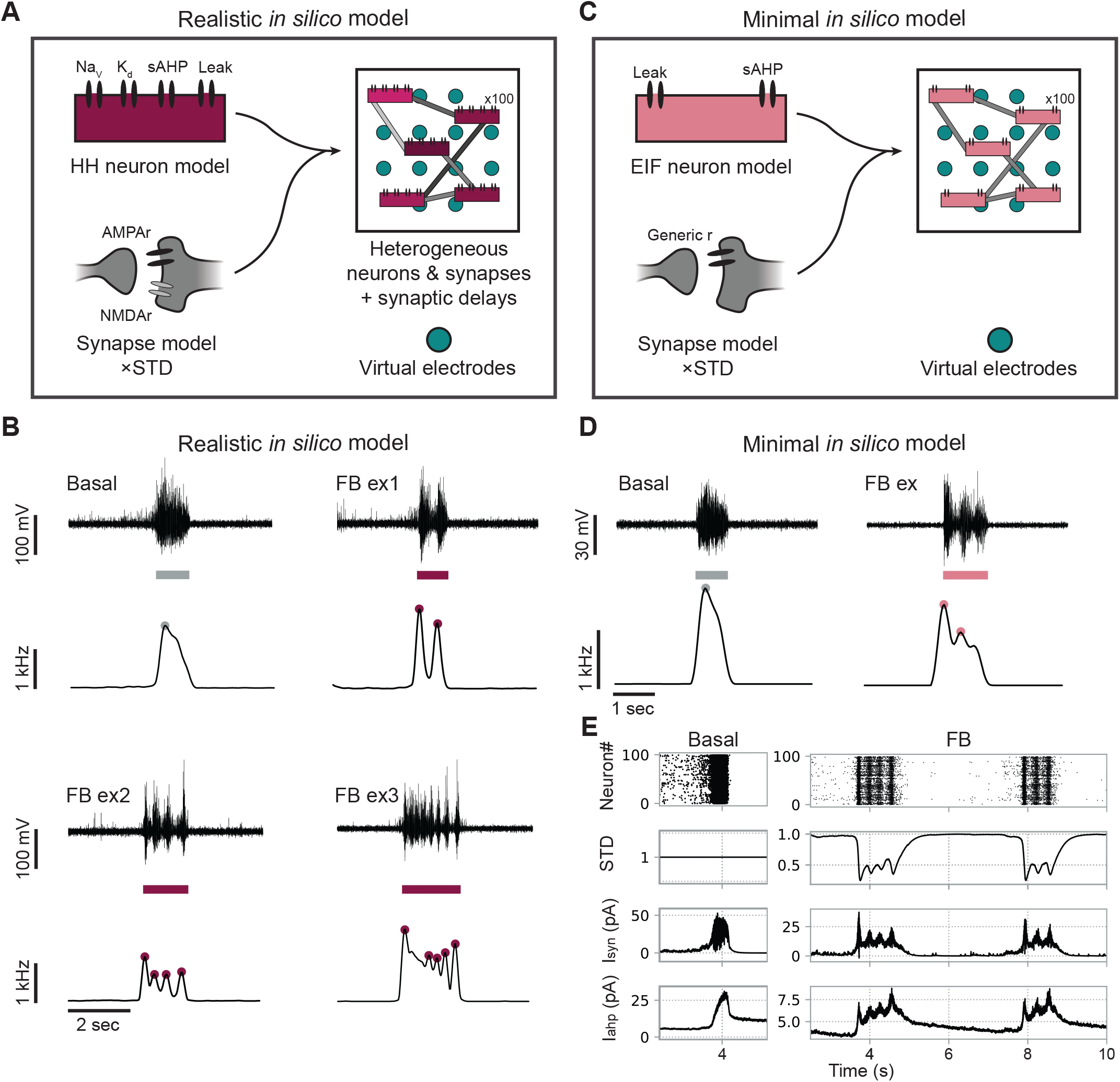
Fragmented network bursts can be replicated in *in silico* models with sufficiently strong short-term synaptic depression. **A)** Schematic representation of the realistic *in silico* model consisting of 100 Hodgkin-Huxley (HH) type neurons with voltage-gated potassium and sodium channels, slow afterhyperpolarizing (sAHP) and leaky currents, connected with AMPA-receptor (AMPAr) and NMDA-receptor (NMDAr) synapses, including short-term depression (STD). All neurons are heterogeneously excitable, with varying synaptic strengths and delays, forming a two-dimensional network. Virtual electrodes are employed to simulate MEA electrodes. **B)** Simulations with the realistic computational model show typical network bursts (NBs) (“Basal”) or, as STD is increased, fragmented NBs (“FB example ex1”). Subsequently, different kinds of fragmented NBs can be created by altering the sAHP and synapse properties (FB examples ex1-ex3). Top panels show voltages recorded at one virtual electrode during a detected NB (colored bar below); bottom panels show the corresponding network firing rate with detected fragments (colored dots). **C)** Minimal *in silico* model with 100 homogeneous exponential integrate-and-fire (EIF) neurons including the sAHP current, homogeneously connected through synapses with generic post-synaptic receptors and STD. **D)** Simulations with the minimal model can also show typical and fragmented NBs. **E)** Bursting mechanism resulting in basal or fragmented NBs. Recurrent excitation (I_syn_, third panel) starts a burst causing rapid firing (top panel). This causes the sAHP current (I_ahp_, fourth panel) to increase, hyperpolarizing the neurons and eventually terminating the NB. In fragmented NBs, the rapid firing causes STD (second panel) to kick in and lower the firing. STD then recovers quickly, allowing the remaining excitation to initiate the next fragment until the sAHP current is high enough to terminate the entire NB.

Fragmented NBs can be simulated with this model by increasing the intensity of STD compared to simulations of healthy controls, resembling fragmented NBs observed *in vitro* (Figure 2B “FB ex1” and Figure 1C). The number and shapes of the fragments could subsequently be modified by additional changes in other model parameters, allowing *in silico* reproduction of the divergent *in vitro* phenotypes (Figure 2B “FB ex2” and “FB ex3” and Figure 1C). In particular, decreased sAHP-currents, increased synaptic strengths or time constants, or decreased STD resulted in more, but less defined fragments (Figure S1). Most importantly, sufficient STD (i.e., strong enough to significantly lower the firing activity) was crucial for the emergence of fragmented NB in all simulations.

Since our *in silico* model consists of many intricate and interacting mechanisms, we could not rule out whether mechanisms other than STD might be involved in the generation of fragmented NBs. To this end, we constructed a simpler computational model only containing the mechanisms we hypothesized to be necessary and sufficient for the emergence of NBs and fragments (i.e. sAHP, recurrent excitation, and STD). In this *minimal in silico* model (Figure 2C), neurons are represented by simple exponential integrate-and-fire (EIF) neurons including the sAHP current. Synapses contain generic synaptic receptors that immediately open in response to a presynaptic spike, and undergo STD; further, all neurons and synapses are identical.

Also with this minimal model, we were able to obtain NBs with and without fragments, and we could identify the mechanisms leading to this phenomenon (Figure 1D). In the simulations, NBs without fragments (Figure 1E, “Basal”) result from the interplay between recurrent excitation (I_syn_) and sAHP (I_AHP_), which suppresses the activity of the neurons. Excitation starts the NB, and the rapid firing causes an increase in the sAHP current. As soon as sAHP is stronger than excitation, the NB stops. Then, sAHP recovers slowly until it is low enough for excitation to initiate the next NB. In simulations with fragmented NBs (Figure 1E, “FB”), STD is strong enough to depress the synapses when rapid firing starts, thereby lowering the firing rate and halting the increase in sAHP. STD then quickly recovers, allowing excitation to become stronger again, and the next fragment starts. This continues until sAHP overcomes excitation and terminates the entire NB. The number of fragments is determined by how fast sAHP overcomes excitation, with more fragments occurring when excitation is stronger than adaptation for longer periods. Our realistic and minimal computational models thus suggest that sufficient STD induces fragmented NBs.

### Enhancing short-term synaptic depression *in vitro* induces fragmented NBs

Next, we aimed to test the hypothesis predicted by our *in silico* model that increased STD causes fragmented NBs by performing a validation *in vitro*. To this end, we applied Dynasore (i.e., dynamin inhibitor that enhances STD [12]) to healthy neuronal networks that showed regular NBs. Upon applying Dynasore (10 *μ*M), fragmented NBs emerged in 50% of the neuronal networks, with two fragments per NB (Figure 3A,B,E).

**Figure 3:**
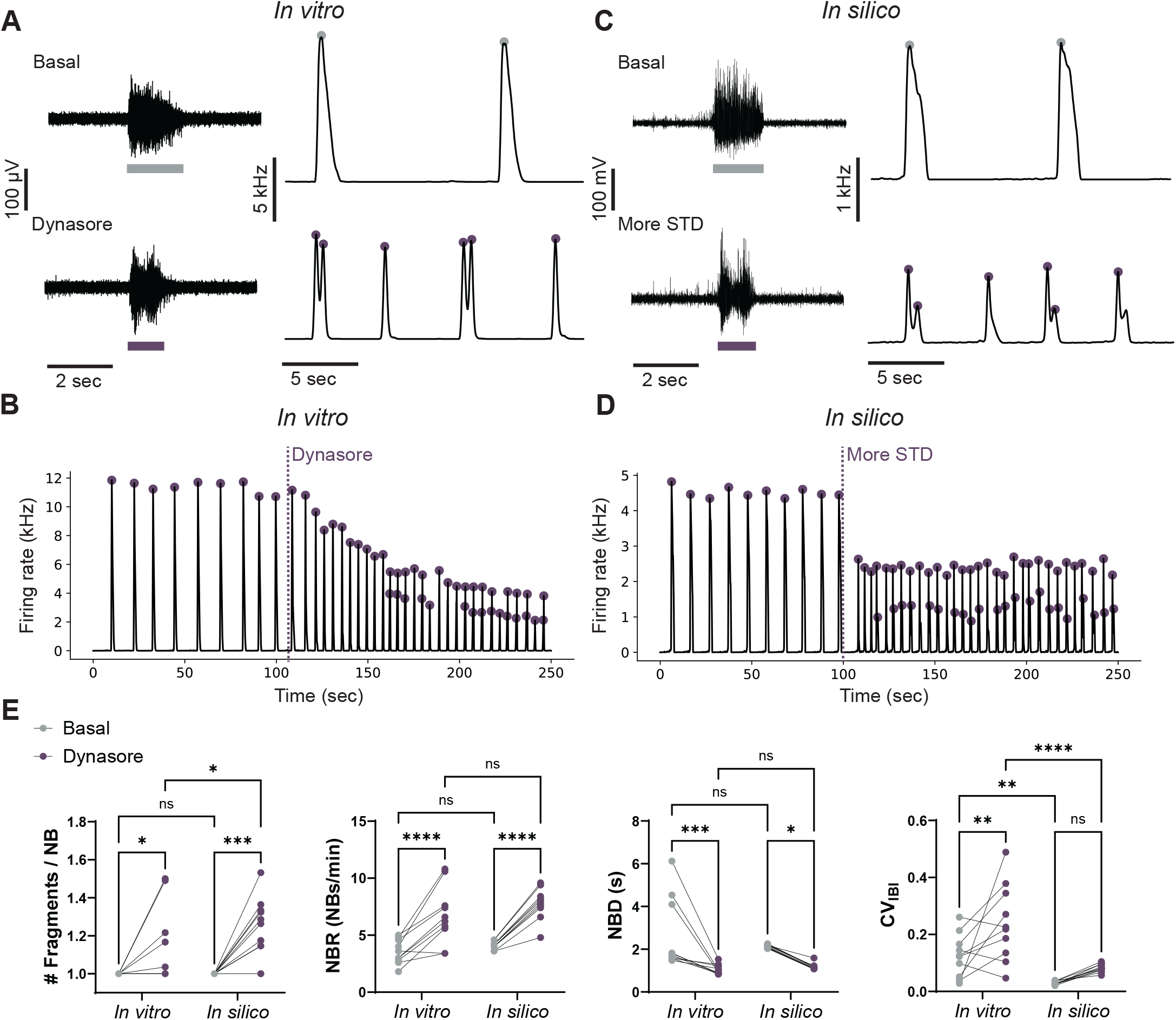
Dynasore induces fragmented network bursts in healthy neuronal networks similar to simulations with increased short-term depression. **A)** Left: example voltage traces recorded at one electrode during a network burst (NB) detection (colored bar below) in a healthy neuronal network *in vitro* before (top) and after (bottom) application of Dynasore (10 *μ*M). Right: example network firing rate traces before (top) and after (bottom) application of Dynasore (detected fragmented are indicated by colored dots). **B)** Representative network firing rate trace when Dynasore is applied to healthy neuronal networks *in vitro* at t=105 s. Purple dots are detected fragments, when two dots appear during one increase in firing rate, two fragments are detected in one NB. **C)** Left: example voltage traces recorded *in silico* by one virtual electrode during a simulation with basal values for short-term depression (STD) (top) and increased values for STD (bottom). Right: example network firing rate traces of simulations with basal values for STD (top) and increased values for STD (bottom). **D)** Representative network firing rate trace when the amount of STD is increased *in silico* at t=100 s. **E)** Quantification of the change when applying Dynasore *in vitro* (n=10) and increasing STD *in silico* (n=10) of the number of fragments per NB (# Fragments / NB), the NB duration (NBD), the NB rate (NBR), and the coefficient of variation of the inter burst intervals (CV_IBI_). ns P*>*0.05, * P*<*0.05, ** P*<*0.01, *** P*<*0.001, **** P*<*0.0001, Two-way ANOVA with uncorrected Fisher’s LSD for multiple comparisons was performed between groups.

To investigate why Dynasore only induced NBs with a maximum of two fragments, we modeled the effect of Dynasore in our realistic *in silico* model by increasing the amount of STD while leaving all other parameters unchanged. Highly similar to the *in vitro* observations, this induced the appearance of either two fragments or of a single fragment with very short NB duration (NBD) (Figure 3C,D,E). We found that when a single fragment occurred, the strength of STD was larger compared to the strength of excitation - which differed per network due to random connectivity - such that sAHP could overcome excitation after the first fragment.

To better compare the *in vitro* effect of Dynasore to the *in silico* increase in STD, we defined additional features of network activity that were affected by Dynasore (Figure 3E). Specifically, the NBD significantly decreased in all wells, and the NB rate (NBR) and coefficient of variation of the inter bursts intervals (CV_IBI_) significantly increased. These changes were not observed when applying a DMSO vehicle to the neuronal networks (Figure S2). Identical to these *in vitro* observations, enhancing STD *in silico* caused an increase in the NBR and a decrease in the NBD. Also, the CV_IBI_ increased *in silico*, albeit not significant, and with notable differences from the *in vitro* observations. To summarize, we show that fragmented NBs can be induced in healthy neuronal networks by enhancing the amount of STD with Dynasore and that the effect of Dynasore was highly similar to enhancing STD *in silico*.

### Persistent excitation in addition to enhanced short-term synaptic depression induces multiple fragments

By only enhancing STD *in vitro*, we could obtain NBs with a maximum of two fragments and a short duration, while patient-derived neuronal networks may show NBs with many more fragments exceeding the NBD of healthy networks (Figure 1C). With our *in silico* models, multiple fragments could be simulated with enhanced STD and excitation stronger than adaptation for a prolonged time (Figure S1). Recently, Pradeepan et al. showed that these multiple fragments occurring in the NBs of neuronal networks derived from Rett syndrome patients *in vitro* (Figure 4A,B) disappeared when blocking asynchronous neurotransmitter release [11]. However, it is unclear how asynchronous neurotransmitter release results in NBs with multiple fragments and whether it is associated with the mechanisms revealed by our *in silico* model.

**Figure 4:**
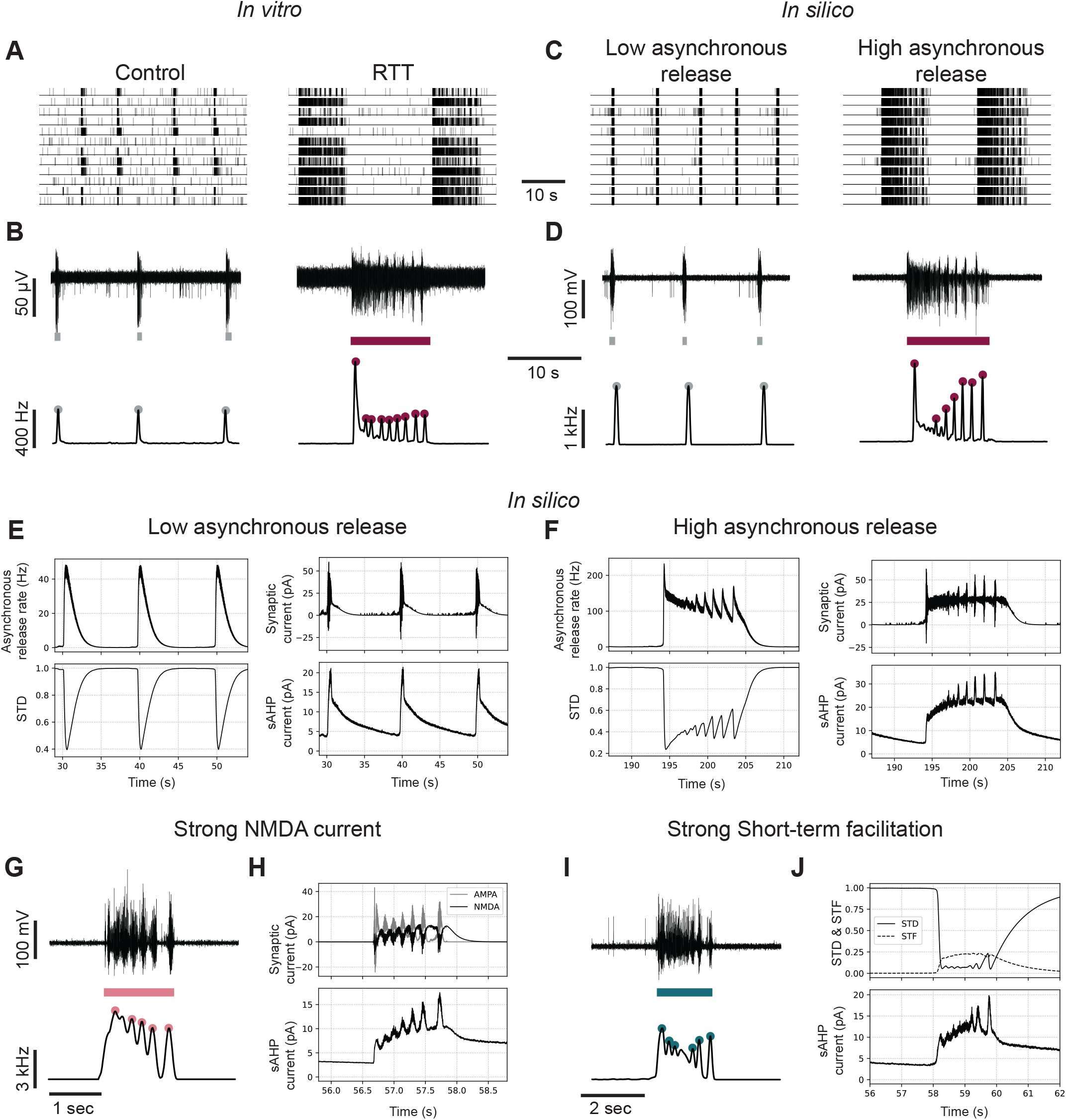
Multiple fragments can be generated *in silico* by enhanced STD with asynchronous release, NMDA currents, or short-term facilitation. **A)** Representative rasterplots of spontaneous neuronal network activity of healthy neuronal networks (left) and neuronal networks derived from a patient with Rett (RTT) syndrome, measured by Pradeepan et al. [11] **B)** Example voltage traces recorded at one electrode in the healthy network (top left) or RTT network (top right) during network burst (NB) detections (colored bar below), and example network firing rate traces during the same period (bottom) showing fragments (colored dots). **C)** Representative rasterplots of simulated activity with low amounts of asynchronous release (left), and high amounts of asynchronous release (right). **D)** Example voltage traces recorded at one virtual electrode in a simulation with low amounts of asynchronous release (top left) or high amounts of asynchronous release (top right) during NB detections (colored bar below), and example network firing rate traces during the same period (bottom) showing fragments (colored dots). **E)** NB mechanism in simulations with low amounts of asynchronous release: the burst ends because there is not enough asynchronous release to initiate the next fragment when sAHP terminates the NB. **F)** NB mechanism in simulations with high amounts of asynchronous release: the firing rate decreases because asynchronous release depletes the amount of available neurotransmitters, but because sAHP is not high enough to terminate the NB and there is enough remaining asynchronous release, the next fragment is initiated, repeating itself until sAHP reaches a threshold. **G)** Top: representative voltage trace recorded at one virtual electrode during a simulation without asynchronous release but with strong NMDA currents and bottom: representative network firing rate trace during the same NB showing multiple fragments. **H)** Mechanism of multiple fragments: a burst fragment is terminated by short-term depression (STD), but because NMDA channels close slowly and only open when the postsynaptic neuron is depolarized, the NMDA current revives the burst causing the next fragment, repeating itself until the sAHP reaches a threshold. **I)** Top: representative voltage trace recorded at one virtual electrode during a simulation without asynchronous release but with short-term facilitation (STF) and bottom: representative network firing rate trace during the same NB showing persistent fragments. **J)** Mechanism of multiple fragments: a burst fragment is terminated by STD, but the burst is revived by STF.

To investigate these mechanisms, we implemented asynchronous transmitter release in our realistic *in silico* model using a phenomenological description [13]. This asynchronous release model describes an increased probability of stochastic neurotransmitter release upon high-frequency pre-synaptic firing, utilizing the same vesicle pool as in “regular” synchronous release. With this model, NBs observed in control neuronal networks could be replicated with low amounts of asynchronous release, while the NBs with multiple fragments observed in diseased networks could be reproduced when the amount of asynchronous release was increased while leaving other parameters unaltered (Figure 4C,D). We could then identify the mechanisms leading to the occurrence of multiple fragments in NBs (Figure 4E,F). In the model, since asynchronous and synchronous release utilize the same vesicle pool, an increase in asynchronous release results in a more depressed synchronous release, and thus increased STD. Hence, fragmentation is still caused by increased STD. However, we noticed that asynchronous release persists long after the pre-synaptic neuron stops firing, allowing the next fragments to be induced after sAHP suppresses the firing. By observing the dynamics of sAHP and excitation, we saw that after every fragment the amount of sAHP is higher and it takes longer before excitation can overcome sAHP to initiate the next fragment, sometimes causing the slowing of the fragments. But, because STD can recover more in this longer period, the synapses will also be stronger with every fragment initiation, causing every subsequent fragment to be more synchronous compared to the last.

Motivated by our hypothesis that multiple fragments are supported by excitation that persists after the pre-synaptic neuron stops firing, we investigated whether also other mechanisms of persistent excitation could support multiple fragment generation. First, since NMDArs open in response to post-synaptic depolarization and typically close slowly, they could cause left-over excitatory currents long after pre-synaptic firing activity stops [14]. We therefore modeled a strong NMDA current in our *in silico* model together with sufficient STD but without asynchronous release. This model indeed also showed NBs with multiple fragments (Figure 4G), where NMDA currents persisted after fragment termination, allowing the next fragment to be initiated (Figure 4H). Short-term synaptic facilitation (STF) can also cause neurotransmitter release after high-frequency firing has ended [15]. To explore this scenario, we constructed a model including STF and sufficient STD but without asynchronous release. In this model, we also observed multiple fragments (Figure 4I,J). Thus, persistent excitation after firing activity ceases, in addition to sufficient STD, supports the generation of multiple fragments.

## Discussion

Fragmented NBs are observed in excitatory neuronal networks derived from patients with several neurological disorders [2, 5, 10, 11]. Different names are given to the phenomenon, such as minibursts, reverberating bursts, high-frequency bursts, and superbursts, but in essence, they are all characterized by a fluctuating spiking frequency within the NBs. In some experimental work, fragmented NBs were suppressed by either NMDAr-blockers [5] or inhibition of asynchronous release [11]. However, none of these studies explain how those processes lead to the formation of fragmented NBs, limiting our understanding of disease mechanisms.

In this work, we aimed to identify candidate biological mechanisms involved in the generation of fragmented NBs, using a biophysical *in silico* model previously developed and validated by us [7], complemented by *in vitro* validations of model predictions.

As confirmed by modeling studies [16, 17, 18, 19], we showed that elementary bursting results from the interplay between excitation and adaptation, modulated by sAHP. Simulations with our biophysical *in silico* model - as well as with our reduced model containing only recurrent excitation, adaptation, and STD - predict that sufficient STD is necessary for the emergence of fragmented NBs. This is similar to previous computational modeling work of primary cultures on MEA, where it was suggested that adaptation, STD, and STF are necessary for the emergence of “repeated network spikes” [18]. However, we show that only STD and adaptation, even without STF, are enough to get fragmentation, dissecting the mechanisms even further. Also, mechanisms proposed in previous work were not verified *in vitro* [18]. Here, we used Dynasore, a drug that enhances STD, to induce fragments *in vitro* and test our hypothesis.

Dynasore inhibits dynamin, a GTPase protein that is essential for endocytosis, and has been shown to inhibit synaptic vesicle recycling in hippocampal neurons in a dose-dependent manner [20]. As a result, Dynasore enhances the amount of STD in these neurons [12]. However, Dynasore might also have other effects on different types of neurons. In brainstem slices, Dynasore completely blocked evoked synaptic responses but increased the spontaneous EPSC (sEPSC) frequency [21]. In the frog neuromuscular junction, Dynasore not only inhibited endocytosis but also led to an increased probability of neurotransmitter release and increased resting intra-terminal calcium. Nevertheless, adding Dynasore in our *in vitro* cultures had an effect that was highly similar to enhancing STD *in silico*, with a maximum of two fragments emerging in some but not all networks, and with identical changes of the NBR and NBD. This suggests that in our hiPSC-derived excitatory cultures, Dynasore works by enhancing the amount of STD. Nevertheless, the effects *in vitro* and *in silico* showed some differences. While the CV_IBI_ significantly increased *in vitro*, the increase *in silico* was not significant, and the values were generally lower. Moreover, the magnitude of the increase in the average number of fragments was higher *in silico* than *in vitro*. Both of these discrepancies might be explained by the fact that the effect of Dynasore changed quite rapidly *in vitro*, while the effect of enhancing STD *in silico* was instant and constant (Figure S3). The effect of Dynasore slowly increased after administration, and then decreased again, causing temporal variations in the number of fragments. At some time points, the effect of Dynasore was so strong that NBDs were shorter than one fragment, and synapses were presumably so depressed that more fragments could not be initiated. Also, IBIs varied with the magnitude of the effect of Dynasore, while this remained constant *in silico*, explaining the higher CV_IBI_ *in vitro*.

In the neuronal networks in which Dynasore caused fragmented NBs, at most two fragments were seen, similar to our *in silico* simulations. This was different from some phenotypes observed *in vitro*, where sometimes up to eight fragments were observed (Figure 1C). While the increase in STD allows the emergence of fragments due to a fast-recovering depression of synaptic transmission, it also allows adaptation to overcome the suppressed excitation faster, terminating the NB sooner. Thus, when only enhancing STD without further altering the excitation-adaptation balance, the NB will always become significantly shorter, and no additional fragments can be generated.

In recent work by Pradeepan et al. [11], multiple fragments occurred in an *in vitro* neuronal network model for Rett Syndrome. These multiple fragments could be blocked using EGTA-AM, suggesting they are caused by asynchronous neurotransmitter release. With our *in silico* models, we showed that the number of fragments in NBs can be increased if excitation dominates over adaptation for a prolonged time. However, how this relates to asynchronous neurotransmitter release is unknown. By implementing asynchronous release in our model, we were able to obtain NBs with multiple fragments in simulations. We showed that asynchronous neurotransmitter release induced NBs with multiple fragments through the same uncovered mechanism since it causes both enhanced STD and prolonged excitation. We were also able to simulate similar NBs using enhanced STD in combination with other mechanisms that prolong excitation, particularly strong NMDA currents, and STF.

NBs with multiple fragments are also observed as a relatively rare phenomenon in healthy neuronal networks dissociated from rodents. [22, 23]. These fragmented NBs are often referred to as reverberations and they usually show a defined shape. The first fragment has a high firing rate and high synchronicity and is followed by a drastic drop in both. Then, for every subsequent fragment, the firing rate and synchronicity increase, as well as the time between fragments [23]. We also observed this slowing and growing shape in our simulations, where it was caused by increasing adaptation and fast-recovering STD.

Through patch-clamp technology, it has been observed that a single neuron in a network can also exhibit reverberating activity when stimulated with a brief pulse [24, 25, 26]. In particular, Lau et al. [24] showed that evoked reverberations could be abolished by EGTA-AM, similar to the model of Rett Syndrome, and that applying Strontium, which elevates asynchronous release, exacerbated reverberations. Moreover, Volman et al. [25] showed, employing a computational network model, that evoked fragments are maintained by enhanced asynchronous transmitter release. Additionally, they showed that a fast time-scale depression is responsible for oscillations during NBs, and a slow time-scale depression is responsible for the termination of the NBs. In our model, sAHP serves as a slow time-scale depression that also terminates the NBs. STD is a fast time-scale depression in our model and indeed causes oscillations within the NB, which we call fragments. This substantiates our hypothesis that STD is still necessary to get fragmentation in NBs. Volman et al. did not include NMDArs in their computational model and argued that, therefore, it was impossible to rule out that NMDA might play a similar role as asynchronous release. Lau et al. found that complete blockage of NMDArs reduced the occurrence and duration of reverberations [24]. Using computational modeling, Wang [27] and Compte et al. [26] found that to obtain reverberations, recurrent excitatory synapses must be dominated by a slow component. With their simulations, they show that network dynamics are essentially asynchronous when NMDArs dominate over AMPArs. Conversely, when AMPArs dominate, the network displays coherent single NBs. Thus, NMDArs are often thought to be important in reverberatory/fragmented activity. Alternatively, Dao Duc et al. [19] used hippocampal cultures and slices, as well as computational modeling, to show that the interplay of STD and STF drives NBs with multiple fragments.

With our biophysical computational model that includes asynchronous release, NMDArs, and STF, we showed that all of those mechanisms allowed the generation of multiple fragments in NBs. However, when using realistic time constants for NMDAr-dynamics and STF, NBs and fragments were generally shorter than in simulations with asynchronous release, suggesting the duration of NBs might be used to differentiate between mechanisms. Yet, the time constants for asynchronous release used here are not based on literature, as it is unknown for these types of cultures. Moreover, we assumed asynchronous and synchronous synaptic release both use the same neurotransmitter vesicle pool, even though these pools are also suggested to be different in some types of neurons [28]. We found that if asynchronous release indeed uses the same vesicle pool, enhanced asynchronous release automatically results in more depressed synchronous release, and thus the emergence of fragments by enhanced STD. In this way, the difference between healthy cultures and Rett cultures from Pradeepan et al. [11] could be modeled solely by increasing the amount of asynchronous release. If both types of release use a different vesicle pool, an additional increase in STD would be needed to simulate the phenotype, as was necessary for simulations of NBs with multiple fragments with NMDArs and STF.

With our results, we can summarize and generalize previous research by showing that NBs with multiple fragments are caused by i) sufficient STD (or another fast-timescale depression that is able to temporarily lower the firing rate) terminating fragments, ii) sufficiently strong excitation (e.g. through asynchronous release, NMDAr or STF) to overcome sAHP and initiate the next fragment, and iii) sAHP (or another slow-timescale depression) to terminate the NB.

We revealed candidate mechanisms that underlie the emergence of fragmented NBs. This can help us understand the driving cause of the phenotypes we observe in patient-derived neuronal networks. In KS networks, fragments could be abolished by applying an NMDAr antagonist, suggesting that enhanced NMDAr function induced the fragmented NBs in those networks [5]. In DS networks, reduced spontaneous sEPSC amplitudes and frequencies were observed, which is hypothesized to be caused by homeostatic synaptic downscaling in response to elevated network activity [7]. Cohen et al. [29] showed that in neuronal networks where the activity was artificially elevated, neurons expressed strong STD that was not seen in untreated neurons. This suggests that homeostatic synaptic downscaling could enhance STD, which might cause the fragmented NBs in DS networks, as well as GEFS+ networks which show a similar phenotype [2]. In both GEFS+ and DS networks, the number of fragments could be increased by elevating the temperature [2]. It has been shown in hippocampal slices that increasing the temperature enhances the amount of STF while leaving the amount of STD unaltered [30]. Thus, at higher temperatures in GEFS+ and DS networks, strong STD due to homeostatic downscaling may remain while STF increases, promoting the initiation of subsequent fragments like in our simulations of NBs with multiple fragments. Moreover, sAHP has been suggested to be lowered in these networks [7], which would allow the occurrence of multiple fragments in combination with enhanced STD (Figure S1).

To conclude, using *in silico* computational models and *in vitro* experiments, we show that enhanced STD is essential for the emergence of fragmented NBs. Asynchronous neurotransmitter release acts on STD, causing fragmentation, but it moreover causes left-over excitation after NB termination, allowing the initiation of many more fragments. These multiple fragments could also be induced with sufficiently strong NMDAr-currents or STF, in combination with sufficient STD.

## Experimental procedures

### *In silico* modelling

All simulations were performed with the Brian2 simulator [31] in a Python 3.9 environment. Differential equations were integrated using either the Exponential Euler or Euler Forward method. The python code to run simulations is published on GitLab.

### Realistic *in silico* model

The realistic *in silico* model is described in Doorn et al. [7]. In short, it consists of one hundred HH-type neurons with voltage-gated sodium and potassium channels, as well as leaky channels and sAHP channels, modeled as a potassium channel whose conductance increases upon action potential (AP) firing. The neurons are heterogeneously excitable through a variable external input current, accounting for intrinsic differences. Additionally, the neurons receive noisy fluctuations of their membrane potential to mimic the effect of sEPSCs or other noise components. Neurons are connected to a subset of other neurons through synapses with models of AMPArs, which open immediately upon arrival of a pre-synaptic spike and decay rapidly, and NMDArs, which open and close slowly and are blocked by magnesium ions that are removed upon depolarization of the post-synaptic neuron. The strengths of the synapses vary due to heterogeneous weights (*w*), drawn from a normal distribution. These weights are further modulated by STD, following the Markram-Tsodysk model [32]. The model is based on the concept of synaptic resources, of which only a fraction, *x*, is available, and of which the release probability (*u*(*t*)) increases upon every pre-synaptic spike. In our original model, we keep *u*(*t*) at a value of 1 to only model STD and not STF. The synaptic weight *w*_*j*_ is multiplied by *x*_*j*_, where *x*_*j*_ obeys:

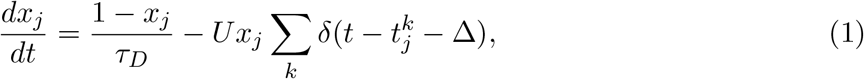

where *τ*_*D*_ is the time constant of STD, and U is the strength of STD.

The parameter values of the model used for all simulations can be found in the Supplemental Table S1.

### Minimal *in silico* model

The minimal model consists of one hundred EIF neurons, where the membrane potential is described by:

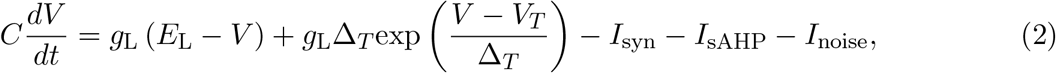

with the reset condition that if *V > V*_*thres*_ then *V* = *V*_*reset*_. *V* is the membrane potential, *C* is the membrane capacitance, *E*_*L*_ is the reversal potential of the leak current and *g*_*L*_ is the corresponding conductance. *I*_*sAHP*_ and *I*_*noise*_ are identical to the realistic model [7] and the synaptic current *I*_*syn*_ is given by:

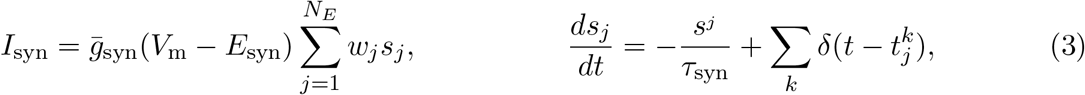

where 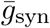 is the maximal synaptic conductance when synaptic channels are opened, *E*_syn_ is the synaptic reversal potential, *w*_*j*_ is the same as in the realistic model and *s* is the fraction of open channels that increases upon every pre-synaptic spike at 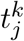 and then decays with time constant *τ*_syn_. The neurons in the model are all identical, as well as the synapses. These synapses also undergo STD as described above. All parameter values used for simulations with the minimal model can be found in the provided python code.

### Short-term facilitation and asynchronous release

To model STF in the model, we use the release probability *u*(*t*) as described by Markram and Tsodysk [32], which, together with the above-mentioned model for STD, forms the short-term plasticity (STP) model:

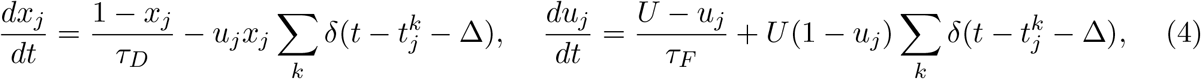

where *τ*_*F*_ is the recovery time constant of facilitation. The synaptic weight is multiplied by both *u*_*j*_ and *x*_*j*_. Thus, at every pre-synaptic spike, *u*_*j*_ increases with *U* (1 − *u*), and an amount of *u*_*j*_*x*_*j*_ neurotransmitters is released and subtracted from *x*_*j*_.

To model asynchronous release, we used an extension of the STP model as described in Wang et al. [13]. In this extension, *u*_*sr*_ describes the release probability of synchronous release as described in equation 4, and *u*_*ar*_ is the probability rate of asynchronous release. Research has shown that synchronous and asynchronous release are mediated by different Ca^2+^ sensors with distinct association and dissociation rates with Ca^2+^ [33]. Therefore, both releases have different recovery time constants (dissociation rates), *τ*_*sr*_ and *τ*_*ar*_, and different saturation levels *U*_*sr*_ and *U*_*ar*_. Based on the literature, we also assume that synchronous and asynchronous releases are competing for the same vesicle pool [33]. We thus use one variable *x* as in equation 4. While synchronous release is time-locked to the arrival of a pre-synaptic spike, asynchronous release is very stochastic. We therefore model it using a binomial process. We call *x*_0_ the amount of neurotransmitter in one vesicle, making *x*(*t*)*/x*_0_ the maximum number of releasable vesicles. In a time-interval [*t, t* + *dt*], the release probability of a single vesicle is given by *u*_*ar*_*dt*. We assume the amount of asynchronous release events *n*(*t*) follows a binomial distribution ℬ (⌊*x*(*t*)*/x*_0_⌋, *u*_ar_(*t*)*dt*). The overall asynchronous release rate is then:

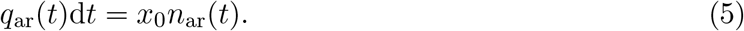

Because we cannot model a binomial distribution in brian2, we approximate it with a normal distribution cut-off at the borders of the binomial distribution. The parameters used for all simulations can be found in the Supplemental Table S1.

### MEA recordings and data analysis

We used MEA recordings performed on neuronal networks derived from hiPSCs of control and patients with GEFS+ and Dravet syndrome (FAM001 GEFS and FAM001 DRAV described in [2]) and control and a patient with KS syndrome (C_MOS_ and KS_MOS_ described in [5]) (data in Figure 1B,C). In addition, we used a representative MEA recording performed on neuronal networks derived from hiPSCs of control and patients with RTT syndrome (described in [11]) (data in Figure 4A,B). To evaluate the effect of enhanced STD *in vitro*, we performed experiments in which we differentiated neuronal networks from hiPSCs of a healthy individual [1] (data in Figure 3). We performed recordings using the Multiwell-MEA system on DIV 35 (Multichannel Systems, MCS GmbH, Reutlingen, Germany). MEA devices are composed of 24 independent wells with embedded micro-electrodes (i.e., 12 electrodes/well, 80 μm in diameter and spaced 300 μm apart). During recording, the temperature was maintained at 37 °C, and a slow flow of humidified gas (5% CO_2_ and 95% ambient air) was applied onto the MEA plate. Every electrode recorded voltages with a sampling frequency of 10 kHz. Also from the *in silico* model, ‘virtual electrode’ recordings were sampled at 10 kHz and then handled identically to experimental data. Wells that did not show electrical activity, that did not show NBs, or that showed unevenly distributed neurons under the microscope were excluded [1].

Signals were filtered between 100 and 3500 Hz using a fifth-order Butterworth filter. We detected spikes using an amplitude threshold-based method, where the threshold was four times the root mean square of the electrode signal. The network firing rate was computed by binning spikes at all electrodes in 25 ms time bins. We then smoothed this network firing rate by convolution with a gaussian kernel. To detect NBs, we employed two thresholds on this smoothed firing rate to start and stop the NB, set to 1/4th and 1/100th of the maximum firing rate respectively. Additionally, 30% of the active electrodes (i.e., electrodes with an average firing rate above 0.02 Hz) had to be firing during the NB, and the firing rate should remain above the NB start- or stop-threshold for 50 ms to start or end NB detection. Fragments were detected using a peak detection algorithm on the smoothed network firing rate where peaks should have a minimal height of 1/16th of the maximum firing rate and a minimal prominence of 1/10th of the maximum firing rate. The number of fragments per NB was then defined as the number of detected peaks within the duration of the NB.

We defined four features that were representative of the spontaneous network activity. Specifically, the average number of fragments per NB (#Fragments/NB) was calculated by summing all detected fragments and dividing them by the number of detected NBs. Additionally, the NBR (the average number of NBs per minute), NBD (the mean duration of NBs during recording), and CV_IBI_ (the coefficient of variation of the inter-burst-intervals) were calculated.

### Dynasore experiments and modeling

Mossink et al. generously provided human induced pluripotent stem cells (hiPSCs), derived from fibroblasts of a healthy individual [1]. We differentiated hiPSCs into excitatory cortical Layer 2/3 neurons on MEA through doxycycline-inducible overexpression of *Neurogenin 2* (*Ngn2*) as described previously [34, 1] (Figure 1A). In short, Ngn2-positive hiPSCs were cultured on Geltrex (Thermofischer) in E8 flex (Gibco), supplemented with puromycin (0.5 μg/ml, Sigma Aldrich), and G418 (50 μg/ml, Sigma Aldrich) at 37°C/5% CO_2_. On DIV0, Ngn2-positive hiPSCs were co-plated as single cells in 24-well MEAs, pre-coated with poly-l-ornithine (50 μg/ml, Sigma Aldrich) and human laminin (5 μg/ml, LN521 BioLamina). To promote neuronal maturation, astrocytes obtained from cortices of newborn (P1) Wistar rats were added to hiPSC cultures in a 1:1 ratio on DIV2. On DIV3, the medium was changed to Neurobasal (Gibco) supplemented with DOX (4 μg/ml, Sigma Aldrich), B-27 (Thermo Fisher Scientific), glutaMAX (2 mM, Thermo Fisher Scientific), primocin (0.1 μg/ml, Invivogen), neurotrophin-3 (10 ng/ml, Bioconnect), and brain-derived neurotrophic factor (10 ng/ml, Bioconnect). Cytosine *β*-D-arabinofuranoside (2 μM, Sigma Aldrich) was added to eliminate proliferating cells. Starting from this day, half of the medium was changed three times a week. From DIV10 onward, 2.5% fetal bovine serum (Sigma Aldrich) was added to support astrocyte viability. DOX was removed after DIV14. Cells were maintained in an incubator at 37°C with 80% humidity and 5% CO_2_ until the experiment on DIV35.

Culturing and experiments were performed with two independent neuronal preparations. Baseline spontaneous activity was recorded for 5 minutes. Immediately following baseline recording, a pharmacological agent (10 μM Dynasore (Sigma Aldrich #1202867-00-2 diluted in DMSO) or 0.1% DMSO) was added to each well, and 20 minutes of post-treatment activity was recorded. Only the last 5 minutes of recording post-treatment were used for analysis.

In the computational model, the effect of Dynasore was modeled by simulating a network for 5 minutes, then increasing *U* from 0.006 to 0.035, while keeping all other parameters identical, and then continuing the simulation for 5 minutes.

### Statistics

We performed statistical analysis using GraphPad Prism 5 (GraphPad Software, Inc., CA, USA). We checked for normal distributions using a Kolmogorov-Smirnov test. We compared the number of fragments in NBs of patient-derived neuronal networks using a Kruskal-Wallis test with Dunn’s multiple comparisons test, because not all data was normally distributed. To compare the effect of Dynasore to vehicle, and to its *in silico* counterpart, we used a two-way ANOVA with uncorrected Fisher’s LSD for multiple comparisons. P-values*<*0.05 were considered significant in all cases. All data points, statistics, and p-values can be found in supplemental table S2.

## Author Contributions

N.D. developed the *in silico* models and designed and performed *in silico* experiments. N.D., E.J.H.F.V., and M.L. designed, prepared and performed *in vitro* experiments. N.D. designed and performed data analyses. N.D. wrote the manuscript with input from all authors. M.F. and M.J.A.M.vP. provided conceptualization and intellectual content. M.F. conceived and supervised the project.

## Acknowledgements

This work was supported by the Netherlands Organisation for Health Research and Development ZonMw BRAINmodel PSIDER program 10250022110003 (to M.F.). We thank Eline van Hugte, Nael Nadif Kasri, and James Ellis for providing MEA recordings from patient-derived *in vitro* neuronal networks.

## Conflict of interest

The authors declare no competing interests.

## Notes

https://gitlab.utwente.nl/m7706783/fb_model

### Competing Interest Statement

The authors have declared no competing interest.

## References

[1] B. Mossink et al. “Human neuronal networks on micro-electrode arrays are a highly robust tool to study disease-specific genotype-phenotype correlations in vitro”. In: Stem Cell Reports 16.9 (Sept. 2021), pp. 2182–2196. doi: 10.1016/J.STEMCR.2021.07.001.

[2] E. J. H. Van Hugte et al. “SCN1A-deficient excitatory neuronal networks display mutation-specific phenotypes”. In: Brain 139.4 (2012), pp. 16–17. doi: 10.1093/BRAIN/AWAD245.

[3] T. M. Klein Gunnewiek et al. “Sonlicromanol improves neuronal network dysfunction and transcriptome changes linked to m.3243A¿G heteroplasmy in iPSC-derived neurons”. In: Stem Cell Reports 16.9 (Sept. 2021), pp. 2197–2212. doi: 10.1016/J.STEMCR.2021.07.002.

[4] T. Klein Gunnewiek et al. “Mitochondrial dysfunction impairs human neuronal development and reduces neuronal network activity and synchronicity”. In: Cell reports (2020), p. 720227. doi: 10.1101/720227.

[5] M. Frega et al. “Neuronal network dysfunction in a model for Kleefstra syndrome mediated by enhanced NMDAR signaling”. In: Nature Communications 10.1 (Oct. 2019), pp. 1–15. doi: 10.1038/s41467-019-12947-3.

[6] M. C. Marchetto et al. “Altered proliferation and networks in neural cells derived from idiopathic autistic individuals”. In: Molecular psychiatry 22.6 (June 2017), pp. 820–835. doi: 10.1038/MP.2016.95.

[7] N. Doorn, E. J. van Hugte, U. Ciptasari, A. Mordelt, H. G. Meijer, D. Schubert, M. Frega, N. Nadif Kasri, and M. J. van Putten. “An in silico and in vitro human neuronal network model reveals cellular mechanisms beyond NaV1.1 underlying Dravet syndrome”. In: Stem Cell Reports 18.8 (Aug. 2023), pp. 1686–1700. doi: 10.1016/J.STEMCR.2023.06.003.

[8] K. Linda et al. “Imbalanced autophagy causes synaptic deficits in a human model for neurodevelopmental disorders”. In: Autophagy 18.2 (2022), pp. 423–442. doi: 10.1080/15548627.2021.1936777.

[9] S. Wang et al. “Loss-of-function variants in the schizophrenia risk gene SETD1A alter neuronal network activity in human neurons through the cAMP/PKA pathway”. In: Cell Reports 39.5 (May 2022), p. 110790. doi: 10.1016/J.CELREP.2022.110790.

[10] M. Gabriele et al. “KMT2D haploinsufficiency in Kabuki syndrome disrupts neuronal function through transcriptional and chromatin rewiring independent of H3K4-monomethylation”. In: bioRxiv (Apr. 2021), p. 2021.04.22.440945. doi: 10.1101/2021.04.22.440945.

[11] K. S. Pradeepan, F. P. McCready, W. Wei, M. Khaki, W. Zhang, M. W. Salter, J. Ellis, and J. Martinez-Trujillo. “Calcium dependent hyperexcitability in human stem cell derived Rett syndrome neuronal networks”. In: Biological Psychiatry Global Open Science (Jan. 2024), p. 100290. doi: 10.1016/J.BPSGOS.2024.100290.

[12] Y. Hua, A. Woehler, M. Kahms, V. Haucke, E. Neher, and J. Klingauf. “Blocking Endocytosis Enhances Short-Term Synaptic Depression under Conditions of Normal Availability of Vesicles”. In: Neuron 80.2 (Oct. 2013), pp. 343–349. doi: 10.1016/J.NEURON.2013.08.010.

[13] T. Wang, L. Yin, X. Zou, Y. Shu, M. J. Rasch, and S. Wu. “A phenomenological synapse model for asynchronous neurotransmitter release”. In: Frontiers in Computational Neuroscience 10.January (Jan. 2016), p. 174362. doi: 10.3389/FNCOM.2015.00153/BIBTEX.

[14] M. Hedegaard, K. B. Hansen, K. T. Andersen, H. Bräuner-Osborne, and S. F. Traynelis. “Molecular pharmacology of human NMDA receptors”. In: Neurochemistry international 61.4 (Sept. 2012), p. 601. doi: 10.1016/J.NEUINT.2011.11.016.

[15] S. L. Jackman and W. G. Regehr. “The Mechanisms and Functions of Synaptic Facilitation”. In: Neuron 94.3 (May 2017), pp. 447–464. doi: 10.1016/J.NEURON.2017.02.047.

[16] M. Augustin, J. Ladenbauer, and K. Obermayer. “How adaptation shapes spike rate oscillations in recurrent neuronal networks”. In: Frontiers in Computational Neuroscience 7.FEB (Feb. 2013), p. 36977. doi: 10.3389/FNCOM.2013.00009/BIBTEX.

[17] A. Compte, M. V. Sanchez-Vives, D. A. McCormick, and X. J. Wang. “Cellular and network mechanisms of slow oscillatory activity (¡1 Hz) and wave propagations in a cortical network model”. In: Journal of Neurophysiology 89.5 (May 2003), pp. 2707–2725. doi: 10.1152/jn.00845.2002.

[18] T. Masquelier and G. Deco. “Network bursting dynamics in excitatory cortical neuron cultures results from the combination of different adaptive mechanisms”. In: PloS one 8.10 (Oct. 2013). doi: 10.1371/JOURNAL.PONE.0075824.

[19] K. Dao Duc, C. Y. Lee, P. Parutto, D. Cohen, M. Segal, N. Rouach, and D. Holcman. “Bursting Reverberation as a Multiscale Neuronal Network Process Driven by Synaptic Depression-Facilitation”. In: PLOS ONE 10.5 (May 2015), e0124694. doi: 10.1371/JOURNAL.PONE.0124694.

[20] J. Newton, T. Kirchhausen, and V. Murthy. “Newton AJ, Kirchhausen T, Murthy VN.. Inhibition of dynamin completely blocks compensatory synaptic vesicle endocytosis. Proc Natl Acad Sci USA 103: 17955-17960”. In: Proceedings of the National Academy of Sciences of the United States of America 103 (Dec. 2006), pp. 17955–17960. doi: 10.1073/pnas.0606212103.

[21] M. E. Hofmann and M. C. Andresen. “Dynasore blocks evoked release while augmenting spontaneous synaptic transmission from primary visceral afferents”. In: PLOS ONE 12.3 (Mar. 2017), e0174915. doi: 10.1371/JOURNAL.PONE.0174915.

[22] D. A. Wagenaar, J. Pine, and S. M. Potter. “An extremely rich repertoire of bursting patterns during the development of cortical cultures”. In: BMC Neuroscience 7.1 (Feb. 2006), pp. 1–18. doi: 10.1186/1471-2202-7-11.

[23] C. H. Huang, Y. T. Huang, C. C. Chen, and C. K. Chan. “Propagation and synchronization of reverberatory bursts in developing cultured networks”. In: Journal of Computational Neuroscience 42.2 (Apr. 2017), pp. 177–185. doi: 10.1007/S10827-016-0634-4.

[24] P. M. Lau and G. Q. Bi. “Synaptic mechanisms of persistent reverberatory activity in neuronal networks”. In: Proceedings of the National Academy of Sciences of the United States of America 102.29 (July 2005), pp. 10333–10338. doi: 10.1073/PNAS.0500717102.

[25] V. Volman, R. C. Gerkin, P. M. Lau, E. Ben-Jacob, and G. Q. Bi. “Calcium and synaptic dynamics underlying reverberatory activity in neuronal networks”. In: Physical Biology 4.2 (June 2007), p. 91. doi: 10.1088/1478-3975/4/2/003.

[26] A. Compte, N. Brunel, P. S. Goldman-Rakic, and X. J. Wang. “Synaptic Mechanisms and Network Dynamics Underlying Spatial Working Memory in a Cortical Network Model”. In: Cerebral Cortex 10.9 (Sept. 2000), pp. 910–923. doi: 10.1093/CERCOR/10.9.910.

[27] X. J. Wang. “Synaptic Basis of Cortical Persistent Activity: the Importance of NMDA Receptors to Working Memory”. In: Journal of Neuroscience 19.21 (Nov. 1999), pp. 9587–9603. doi: 10.1523/JNEUROSCI.19-21-09587.1999.

[28] P. S. Kaeser and W. G. Regehr. “Molecular Mechanisms for Synchronous, Asynchronous, and Spontaneous Neurotransmitter Release”. In: Annual review of physiology 76 (Feb. 2014), p. 333. doi: 10.1146/ANNUREV-PHYSIOL-021113-170338.

[29] D. Cohen and M. Segal. “Homeostatic presynaptic suppression of neuronal network bursts”. In: Journal of neurophysiology 101.4 (Apr. 2009), pp. 2077–2088. doi: 10.1152/JN.91085.2008.

[30] V. A. Klyachko and C. F. Stevens. “Temperature-Dependent Shift of Balance among the Components of Short-Term Plasticity in Hippocampal Synapses”. In: Journal of Neuroscience 26.26 (June 2006), pp. 6945–6957. doi: 10.1523/JNEUROSCI.1382-06.2006.

[31] M. Stimberg, R. Brette, and D. F. Goodman. “Brian 2, an intuitive and efficient neural simulator”. In: eLife 8 (Aug. 2019). doi: 10.7554/eLife.47314.

[32] H. Markram, Y. Wang, and M. Tsodyks. “Differential signaling via the same axon of neocortical pyramidal neurons”. In: Proceedings of the National Academy of Sciences of the United States of America 95.9 (Apr. 1998), pp. 5323–5328. doi: 10.1073/pnas.95.9.5323.

[33] H. Wen, M. W. Linhoff, M. J. McGinley, G. L. Li, G. M. Corson, G. Mandel, and P. Brehm. “Distinct roles for two synaptotagmin isoforms in synchronous and asynchronous transmitter release at zebrafish neuromuscular junction”. In: Proceedings of the National Academy of Sciences of the United States of America 107.31 (Aug. 2010), pp. 13906–13911. doi: 10.1073/PNAS.1008598107.

[34] M. Frega, S. H. Van Gestel, K. Linda, J. Van Der Raadt, J. Keller, J. R. Van Rhijn, D. Schubert, C. A. Albers, and N. N. Kasri. “Rapid neuronal differentiation of induced pluripotent stem cells for measuring network activity on micro-electrode arrays”. In: Journal of Visualized Experiments 2017.119 (Jan. 2017), p. 54900. doi: 10.3791/54900.

