## Supplemental Figures and Tables for "Breaking the Burst: Unveiling Mechanisms Behind Fragmented Network Bursts in Patient-derived Neurons"

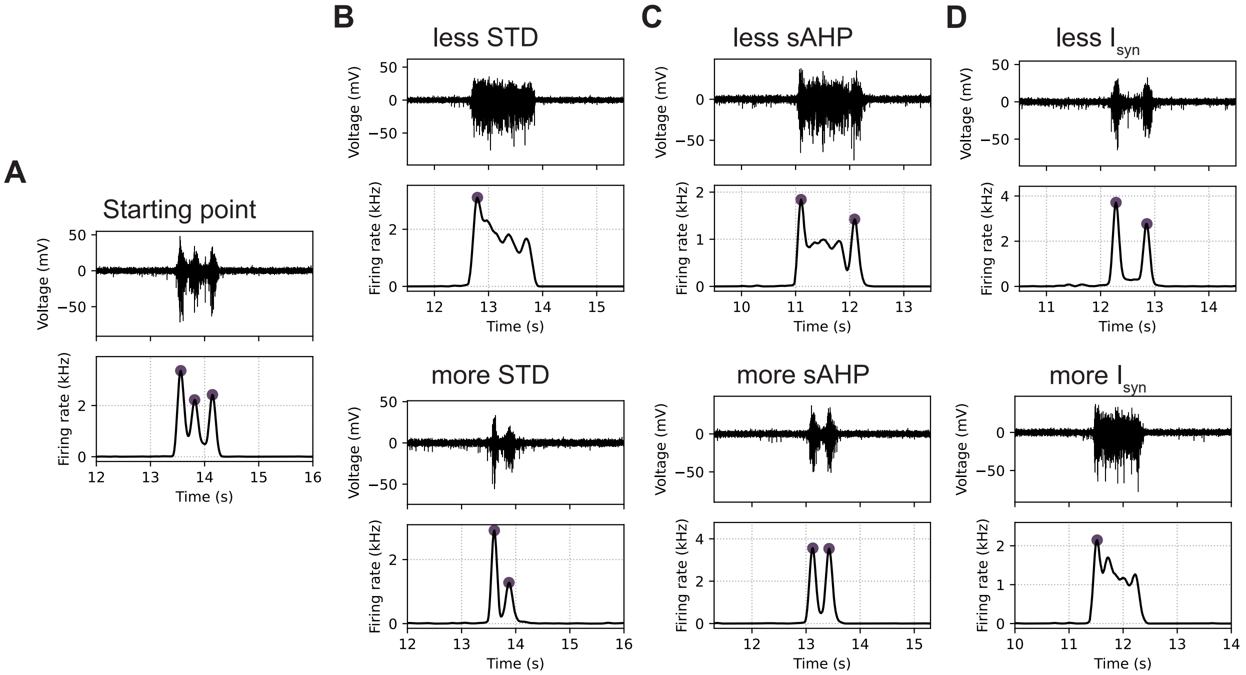


Figure S1: **The effect of model parameters on the number and shape of fragments in simulations. A)** Starting point of parameter configuration resulting in simulations with network bursts (NBs) consisting of three fragments. **B)** The effect of less or more STD on the simulated NBs. Modeled as an increase or decrease in parameter $U$. **C)** The effect of less or more sAHP current on the simulated NBs. Modeled as an increase or decrease in the conductance of the sAHP channels. **D)** The effect of less or more synaptic current ($I_{syn}$) on the simulated NBs. Modeled as an increase or decrease in the conductance of all synaptic channels.

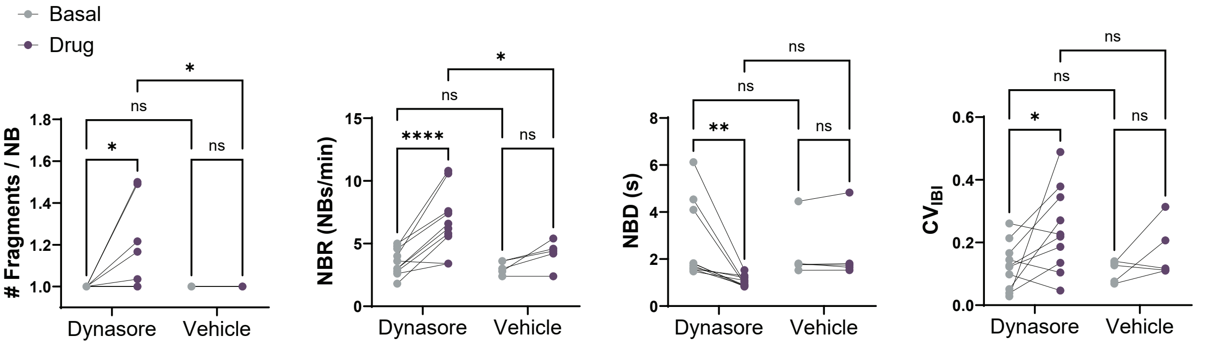


Figure S2: **Dynasore affects neuronal network activity differently from vehicle.** The effect on the neuronal network activity when either Dynasore (10 $\mu$M) (n=10) or vehicle (0.1% DMSO) (n=5) is applied to the network. Figure shows the change in the average number of fragments per network burst (NB) (# Fragments / NB), the NB rate (NBR), the NB duration (NBD), and the coefficient of variation of the inter burst intervals ($CV_{IBI}$). ns P>0.05, * P<0.05, ** P<0.01, **** P<0.0001, 2way ANOVA with uncorrected Fisher's LSD for multiple comparisons was performed between groups.

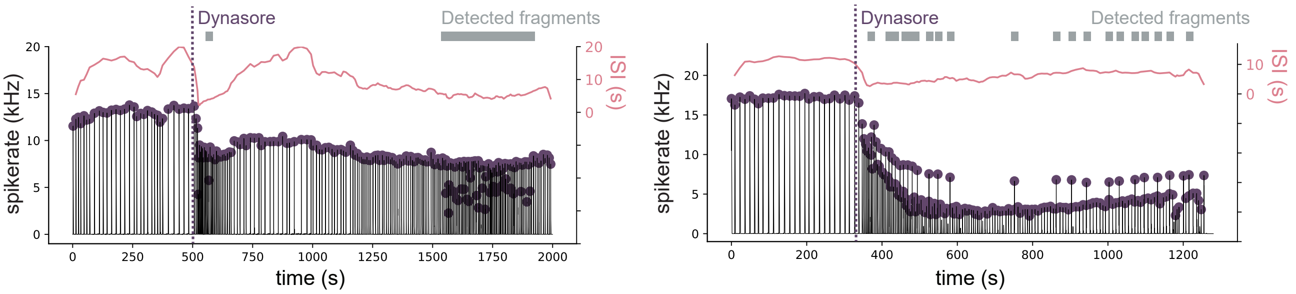


Figure S3: **The effect of Dynasore on the number of fragments and inter-spike interval varies throughout the measurement.** Two examples of recordings of healthy neuronal network activity in which Dynasore was added (dashed line). Shown is the network firing rate in black with detected peaks in purple. In both examples, a decrease in the network firing rate can be seen upon Dynasore application, together with a decrease in the inter-spike interval (ISI, pink line) and some fragment detections (grey bar). Then the ISI slowly increases again while the amount of fragment detections drops to zero. Towards the end of the recording, the amount of fragment detections again increases, illustrating the variable effect Dynasore has on the *in vitro* neuronal network over time.

### Supplemental Tables

Table S1: Values of the parameters of the realistic *in silico* model used for the results and examples shown in the Figures.

| **Parameter** | **Unit** | **Fig 2B** | **Fig 3C,D** | **Fig 4C,D** | **Fig 4G,H** | **Fig 4I,J** |
|---|---|---|---|---|---|---|
| Connection prob | | 0.2 | 0.2 | 0.2 | 0.3 | 0.2 |
| Membrane area | $\mu m^{-2}$ | 300 | 300 | 300 | 300 | 300 |
| **C** | $\mu F \cdot cm^{-2}$ | 2 | 2 | 2 | 1 | 2 |
| $E_L$ | mV | -39.2 | -39.2 | -39.2 | -39.2 | -39.2 |
| $E_K$ | mV | -80 | -80 | -80 | -80 | -80 |
| $E_{Na}$ | mV | 70 | 70 | 70 | 70 | 70 |
| $g_{Na}$ | $mS \cdot cm^{-2}$ | 80 | 80 | 80 | 50 | 80 |
| $g_K$ | $mS \cdot cm^{-2}$ | 6.5 | 6.5 | 6.5 | 5 | 6.5 |
| $g_L$ | $mS \cdot cm^{-2}$ | 0.3 | 0.3 | 0.3 | 0.3 | 0.3 |
| Firing threshold | mV | -30.4 | -30.4 | -30.4 | -30.4 | -30.4 |
| $\sigma$ | mV | 8 | 6 | 5.5 | 3.5 | 5.5 |
| $g_{sAHP}$ | nS | 5 | 4 | 8 | 5 | 5 |
| $\tau_{AHP}$ | ms | 8000 | 8000 | 8000 | 8000 | 8000 |
| $\alpha_{Ca}$ | | 0.00035 | 0.00035 | 0.00035 | 0.00035 | 0.00035 |
| S | | 0.4 | 0.25 | 0.4 | 1.5 | 13 |
| $\delta$ | | 0.6 | 0.6 | 0.6 | 0.5 | 0.6 |
| $E_{AMPA}$ | mV | 0 | 0 | 0 | 0 | 0 |
| $E_{NMDA}$ | mV | 0 | 0 | 0 | 0 | 0 |
| $\tau_{AMPA}$ | ms | 2 | 2 | 2 | 2 | 2 |
| $\tau_{NMDA,rise}$ | ms | 2 | 2 | 2 | 2 | 2 |
| $\tau_{NMDA,decay}$ | Ms | 100 | 100 | 100 | 100 | 100 |
| $\alpha$ | kHz | 0.5 | 0.5 | 0.5 | 0.5 | 0.5 |
| $\tau_D$ | ms | 200 | 250 | 200 | 800 | 1000 |
| $U$ | | 0.2 | 6e-3/35e-3 | 0.01 | 0.2 | 0.005 |
| Apply STF | t/f | FALSE | FALSE | FALSE | FALSE | TRUE |
| $\tau_F$ | ms | - | - | - | - | 1000 |
| Apply Asyn | t/f | FALSE | FALSE | TRUE | FALSE | FALSE |
| $\tau_{ar}$ | ms | - | - | 700 | - | - |
| $U_{ar}$ | | - | - | 0.5 | - | - |
| $U_{max}$ | 1/ms | - | - | 5e-4/45e-4 | - | - |
| $X_0$ | | - | - | 5 | - | - |
